# Evolutionary conservation of midline repulsive signaling by Robo family receptors in flies and mice

**DOI:** 10.1101/2025.03.07.642111

**Authors:** Allison Loy, Trent Daiber, Timothy A. Evans

## Abstract

The Roundabout (Robo) family is an evolutionarily conserved group of axon guidance receptors that regulate midline crossing in a wide range of animal groups by signaling midline repulsion in response to Slit ligands. However, it is not known whether Robo receptors from different species signal midline repulsion via equivalent mechanisms. To examine the evolutionary conservation of midline repulsive signaling, here we express Robo family proteins from mouse in neurons of the fruit fly, *Drosophila melanogaster*. We show that Robo proteins which normally function in canonical Slit-dependent midline repulsion in the mouse (mRobo1 and mRobo2) can also repel *Drosophila* axons from the midline in the fly embryonic ventral nerve cord, and can partially rescue the ectopic midline crossing defects caused by loss of *Drosophila robo1*, though less efficiently as *Drosophila* Robo1 itself. In contrast, mouse Robo3 isoforms (mRobo3.1 and mRobo3.2) which do not normally respond to Slit have no detectable effect on axon guidance when expressed in fly embryos. We further show that the differences in midline repulsive signaling effectiveness between fly and mouse Robos is not due to a simple difference in Slit affinity conferred by the Slit-binding Ig1 domain. Together, our results support the idea that the core signaling mechanisms employed by Robo family receptors are conserved across bilaterians, though some degree of evolutionary divergence has decreased the ability of receptors from different bilaterian clades to directly substitute for one another.

## Introduction

The Roundabout (Robo) family of axon guidance receptor proteins is evolutionarily conserved among bilaterian animals, and the canonical role of Robo family members is to signal midline repulsion of axons in response to their cognate Slit ligands. The family was first identified in *Drosophila*, where mutations in the founding member *robo1* cause ectopic midline crossing of axons in the embryonic ventral nerve cord (Seeger et al. 1993). Soon thereafter, Robo homologs were identified in other animal genomes, including mammals and nematode worms, demonstrating the family’s ancient origins and wide evolutionary conservation (Kidd et al. 1998; Zallen et al. 1998; Brose et al. 1999). Most Robo family members exhibit a common structural arrangement of five immunoglobulin-like (Ig) domains combined with three fibronectin-like (Fn) repeats in their extracellular portion, followed by a single transmembrane domain and a cytoplasmic region containing a number of short conserved cytoplasmic (CC) sequence motifs that were identified by virtue of their similarities in the cytoplasmic domains of *Drosophila* and human Robo orthologs (Kidd et al. 1998; Bashaw et al. 2000). Biochemical studies revealed that cross-species interactions of ligand and receptor were possible, with insect Slit proteins binding to mammalian Robo proteins, and vice versa (Brose et al. 1999). This, together with the conservation of cytoplasmic CC motifs, suggested that the signaling mechanisms which activate Robo receptors and induce axon repulsion might be similarly well conserved.

In *Drosophila*, three Robo receptors are present (Robo1, Robo2, and Robo3), and Robo1 is the main midline repulsive receptor and accounts for most of Slit-dependent midline repulsion (Kidd et al. 1998; Brose et al. 1999; Rajagopalan, Nicolas, et al. 2000; Simpson, Kidd, et al. 2000). Robo2 also contributes to Slit-dependent midline repulsion, and *robo1, robo2* double mutants phenocopy *slit* mutants (Rajagopalan, Nicolas, et al. 2000; Simpson, Kidd, et al. 2000). Robo2 also plays additional roles in the embryonic CNS, including promoting midline crossing by antagonizing Slit-Robo1 repulsion in trans (Spitzweck et al. 2010; Evans et al. 2015), guiding longitudinal axons to the lateral regions of the neuropile (Rajagopalan, Vivancos, et al. 2000; Simpson, Bland, et al. 2000; Evans and Bashaw 2010; Spitzweck et al. 2010), and guiding a subset of motor axons to their target muscles in the body wall (Santiago et al. 2014). Robo3 does not appear to contribute to midline repulsion in the embryonic CNS, though it can repel axons in response to Slit and some of its roles do appear to be Slit-dependent (Zlatic et al. 2003; Pappu et al. 2011); instead its primary role in the embryonic ventral nerve cord is to position longitudinal axons within intermediate regions of the neuropile (Rajagopalan, Vivancos, et al. 2000; Simpson, Bland, et al. 2000; Spitzweck et al. 2010), a role that appears to be Slit-independent (Carranza et al. 2023).

In the mouse spinal cord, two Robo paralogs (mouse Robo1 [hereafter mRobo1], and mouse Robo2 [mRobo2]) cooperate to signal midline repulsion in response to three Slit paralogs (mSlit1, mSlit2, and mSlit3) expressed at the spinal cord floor plate (Kidd et al. 1998; Brose et al. 1999; Long et al. 2004; Jaworski et al. 2010). mRobo1 and mRobo2 also regulate longitudinal guidance of post-crossing commissural axons, though whether this activity is equivalent to that of *Drosophila* Robo2 and Robo3 is not yet known (Jaworski et al. 2010). A third mouse Robo paralog (mRobo3, also known as Rig-1) is reported to be alternatively spliced, with its two isoforms (mRobo3.1 and mRobo3.2, which differ at a single small C-terminal exon) playing different roles with respect to midline crossing (Sabatier et al. 2004; Chen et al. 2008). Mammalian Robo3 homologs have lost the ability to bind Slit via a number of acquired amino acid changes in the Ig1 domain (Zelina et al. 2014), and instead respond to the NELL2 ligand to repel axons from lateral regions of the spinal cord and toward the floor plate midline (Jaworski et al. 2015; Pak et al. 2020). Thus, some roles of Robo receptors (in particular, Slit-dependent midline repulsion) appear to be evolutionarily conserved, while others may be unique to specific taxa. It is not yet clear whether the specific mechanisms by which Robo homologs in various species signal midline repulsion in response to Slit are also evolutionarily conserved.

Some trans-species experiments have examined the conservation of Robo receptor midline repulsive signaling by expressing Robo proteins from other species in *Drosophila* neurons. For example, Robo proteins from the flour beetle *Tribolium castaneum* and nematode worm *Caenorhabditis elegans* can activate midline repulsion in *Drosophila* neurons, and can even rescue midline repulsion defects in *robo1* mutant embryos to varying degrees (Evans and Bashaw 2012; Daiber et al. 2021). Similar experiments with mammalian Robo proteins have produced opposite results: misexpression of human Robo1 in *Drosophila* neurons appears to interfere with midline repulsion, inducing a dominant negative-like ectopic midline crossing phenotype, perhaps suggesting that human Robo receptors are unable to recognize or respond to *Drosophila* Slit in vivo (Justice et al. 2017). However, there is precedence for human axon guidance receptors rescuing their orthologous *Drosophila* mutants; for example human Deleted in Colorectal Cancer (DCC) can substitute for its *Drosophila* ortholog Frazzled (Fra) to promote midline attraction of axons in the fly embryonic ventral nerve cord (O’Donnell and Bashaw 2013).

Here, we examine the evolutionary conservation of midline repulsive signaling between Robo family proteins in flies and mice by expressing mouse Robo receptors in fly neurons during embryonic development, and assaying their ability to induce ectopic midline repulsion and to rescue *robo1* mutant ectopic midline crossing phenotypes. We show that the two mouse Robo receptors that normally function in canonical Slit-dependent midline repulsion in the murine spinal cord (mRobo1 and mRobo2) can also repel *Drosophila* axons from the fly embryonic ventral nerve cord midline, and can partially rescue the ectopic midline crossing defects caused by loss of *Drosophila robo1* when expressed at high levels using the GAL4/UAS system, but isoforms of mRobo3/Rig-1, which does not normally participate in Slit-dependent repulsion in mice, cannot. Neither mRobo1 nor mRobo2 can signal midline repulsion in fly neurons as efficiently as *Drosophila* Robo1, as they are unable to rescue *robo1* mutant phenotypes when expressed at *robo1’s* endogenous expression levels. We show that this difference in signaling effectiveness is not due to simple Ig1-dependent differences in affinity for *Drosophila* Slit, as exchanging the Slit binding Ig1 domain in mRobo1 or mRobo2 with the equivalent domain from *Drosophila* Robo1 does not confer Robo1-like levels of midline repulsive activity.

## Materials and Methods

### Molecular biology

#### pUAST cloning

Mouse *robo* coding sequences were amplified via PCR using primers designed against the reported coding sequences from cDNA prepared from embryonic day 15.5 (E15.5) mouse dorsal spinal cord and cloned via CloneJET PCR cloning kit (Thermo Scientific). No recovered clones matched the full reported sequences for *robo3*.*1* or *robo3*.*2*, so the reported sequences were recapitulated by combining PCR fragments from multiple *robo3* clones via Gibson assembly. Verified coding sequences were then cloned via Gibson assembly into a *BglII*-digested pUAST vector (p10UASTattB) including 10xUAS and an attB site for phiC31-directed site-specific integration. *Drosophila robo1* and mouse *robo* p10UASTattB constructs include identical heterologous 5′ UTR and signal sequences (derived from the Drosophila *wingless* gene) and an N-terminal 3xHA tag (Brown et al. 2015). Mouse *robo* constructs include the following amino acid residues after the N-terminal HA tag, relative to the indicated Genbank reference sequences: mRobo1 (XP_006523028.1, P29–S1615); mRobo2 (NP_780758.3, P31– E1508); mRobo3.1 (NP_001158239.1, P64–R1402); mRobo3.2 (EDL25431.1, P42–K1345).

#### robo1 rescue construct cloning

Construction of the *robo1* genomic rescue construct was described previously (Brown et al. 2015). Mouse *robo* coding sequences were cloned via Gibson assembly into the *BamHI*-digested backbone. Receptor proteins produced from this construct include the endogenous Robo1 signal peptide, and the 4xHA tag is inserted directly upstream of the Ig1 domain. For mRobo/dRobo1 Ig1 chimeric receptors, *robo1, mRobo1*, and *mRobo2* receptor fragments were amplified separately via PCR, then assembled and cloned into the *robo1* rescue construct backbone using Gibson Assembly (New England Biolabs E2611). Coding regions for all transgenic constructs were sequenced to ensure no other changes were introduced. Ig1 chimeras include the following amino acid residues following the N-terminal 4xHA tag, relative to the mRobo Genbank sequences listed above and AAF46887.1 (*Drosophila* Robo1): mRobo1^dR1Ig1^ (Robo1 Q52-V152/mRobo1 L129-S1615), mRobo2^dR1Ig1^ (Robo1 Q52-V152/mRobo2 L131-E1508).

### Genetics

The following *Drosophila* mutant alleles were used: *robo1*^*1*^ (also known as *robo*^*GA285*^) (Kidd et al. 1998), *eg*^*Mz360*^ (eg-GAL4) (Dittrich et al. 1997). The following *Drosophila* transgenes were used: *P{GAL4-elav*.*L}3 (elavGAL4)* (Ogienko et al. 2020), *P{UAS-TauMycGFP}III (UAS-TMG), P{10UAS-HA-Robo1}86Fb (UAS-Robo1)* (Evans and Bashaw 2012), *P{10UAS-HA-mRobo1}86Fb (UAS-mRobo1), P{10UAS-HA-mRobo2}86Fb (UAS-mRobo2), P{10UAS-HA-mRobo3*.*1}86Fb (UAS-mRobo3*.*1), P{10UAS-HA-mRobo3*.*2}86Fb (UAS-mRobo3*.*2), P{robo1::HA-robo1}* (Brown et al. 2015), *P{robo1::HA-mRobo1}, P{robo1::HA-mRobo2}, P{robo1::HA-mRobo3*.*1}, P{robo1::HA-mRobo3*.*2}*. Transgenic flies were generated by BestGene Inc (Chino Hills, CA) using ΦC31-directed site-specific integration into attP landing sites at cytological position 86FB (for *UAS-Robo* transgenes) or 28E7 (for *robo1* genomic rescue constructs). *robo1* rescue transgenes were introduced onto a *robo1*^*1*^ chromosome via meiotic recombination, and the presence of the *robo1*^*1*^ mutation was confirmed in all recombinant lines by DNA sequencing. All crosses were carried out at 25°C.

### Immunofluorescence and imaging

*Drosophila* embryo collection, fixation and antibody staining were carried out as previously described (Patel 1994; Hauptman et al. 2022; Kidd and Evans 2023a). The following antibodies were used: FITC-conjugated goat anti-HRP (Jackson Immunoresearch #123-095-021, 1:100); mouse anti-Fasciclin II (Developmental Studies Hybridoma Bank [DSHB] #1D4, 1:100); mouse anti-βgal (DSHB #40-1a, 1:150); rabbit anti-GFP (Invitrogen #A11122, 1:1000); mouse anti-HA (BioLegend #901503, 1:1000); Cy3-conjugated goat anti-mouse (Jackson #115-165-003, 1:1000); Alexa 488–conjugated goat anti-rabbit (Jackson #111-545-003, 1:500). Embryos were genotyped using balancer chromosomes carrying *lacZ* markers, or by the presence of epitope-tagged transgenes. Ventral nerve cords from embryos of the desired genotype and developmental stage were dissected and mounted in 70% glycerol/PBS (Kidd and Evans 2023b). Fluorescent confocal stacks were collected using a Leica SP5 confocal microscope and processed by Fiji/ImageJ (Schindelin et al. 2012) and Adobe Photoshop software.

## Results

### Mammalian Robo proteins can activate midline repulsion in *Drosophila*

To test whether mammalian Robo proteins can influence axon guidance in *Drosophila* neurons, we created *UAS* transgenic lines for mouse Robo1 (mRobo1), mouse Robo2 (mRobo2), and the two reported mouse Robo3/Rig-1 variants (mRobo3.1 and mRobo3.2). We crossed these *UAS-mRobo* lines to *elav-GAL4*, which is expressed broadly in all neurons, and *eg-GAL4*, which is expressed in two subsets of commissural neurons (the EG and EW neurons). We assayed midline axon guidance in these embryos using anti-HRP (which cross-reacts with a pan-neural epitope in *Drosophila* and labels all of the axons in the ventral nerve cord) and anti-FasII (which labels a subset of longitudinal axons) or anti-GFP (which detects TauMycGFP [TMG] expression in EG and EW neurons in embryos carrying *eg-GAL4* and *UAS-TMG)*. As a control for these gain of function experiments, we used a *UAS-Robo1* transgene to misexpress *Drosophila* Robo1 protein using the same GAL4 drivers. Importantly, all of the *UAS* transgenes contained the same 5’UTR, signal sequence, and N-terminal 3xHA epitope tag, and all were inserted at the same genomic location (86FB) to ensure equivalent transcription levels between the transgenes. We also used anti-HA to directly compare transgenic protein expression levels and localization in each experiment.

In *elav-GAL4/UAS-Robo1* embryos, *Drosophila* Robo1 is misexpressed at high levels in all embryonic neurons, resulting in ectopic midline repulsion and a reduction or absence of commissural axon tracts **(Figure 1B)** (Brown et al. 2015). We found that transgenic mRobo1, mRobo2, mRobo3.1, and mRobo3.2 proteins were expressed at similar levels to transgenic *Drosophila* Robo1 when misexpressed with *elav-GAL4*, and all of the mouse Robo proteins were properly localized to axons in the embryonic ventral nerve cord **(Figure 1G-K)**. mRobo1 and mRobo2 were able to induce high levels of ectopic midline repulsion **(Figure 1C,D)**, but mRobo3.1 and mRobo3.2 exhibited no detectable phenotypic effect under equivalent misexpression conditions **(Figure 1E,F)**. We noted that the mRobo2 gain of function phenotype was consistently stronger than mRobo1, with some *elav-GAL4/UAS-mRobo2* embryos exhibiting similar levels of ectopic repulsion to *elav-GAL4/UAS-Robo1*. We also note that mRobo3.1 and mRobo3.2 proteins were present at similar levels on longitudinal and commissural axons, indicating that these two proteins were not downregulated on commissures.

**Figure 1.**
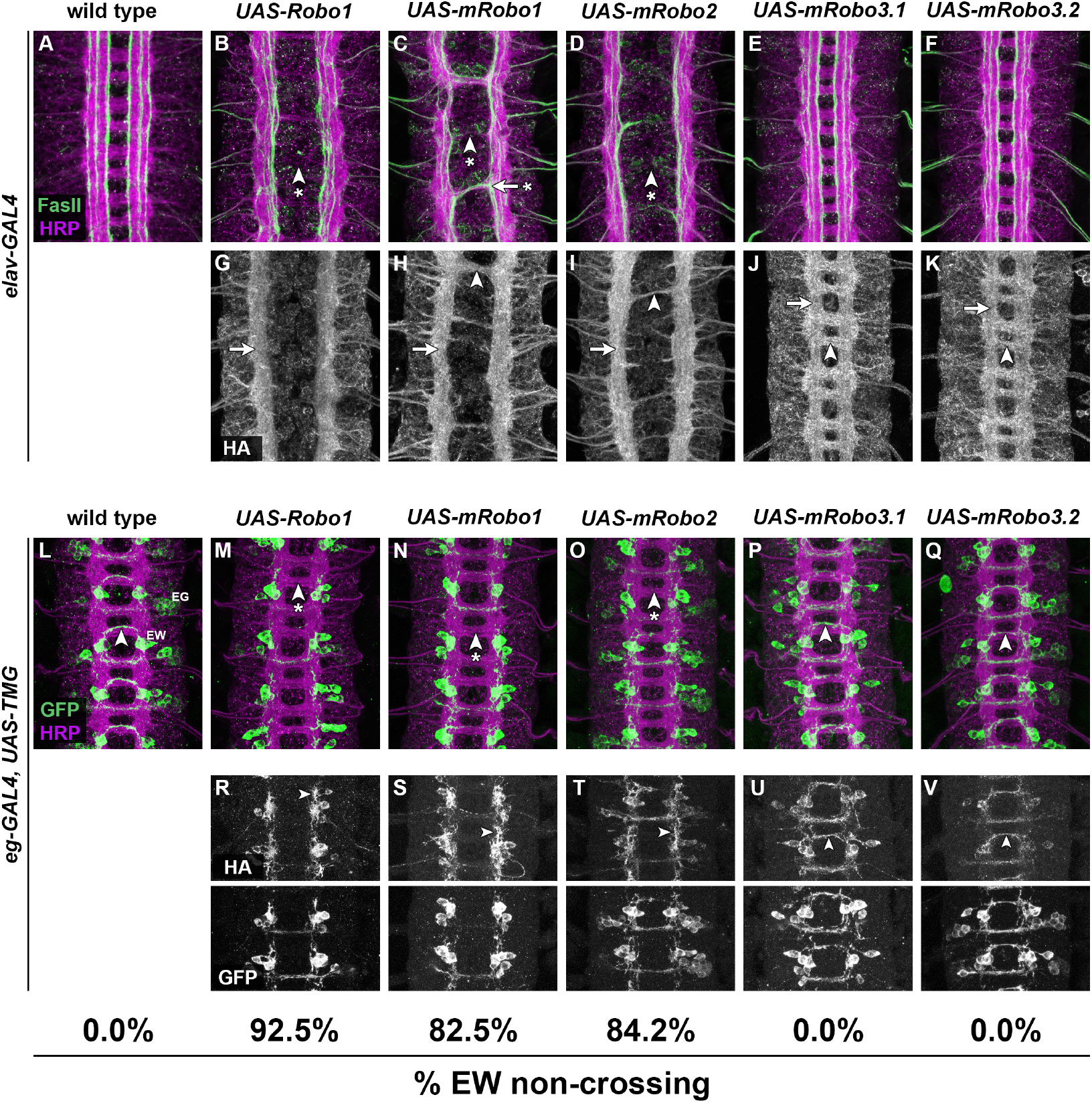
Transgenic mRobo1 and mRobo2 can signal midline repulsion in *Drosophila* neurons. (A-K) Ventral nerve cords from *Drosophila* embryos carrying *elav-GAL4* and the indicated HA-tagged *UAS-Robo* transgenes, stained with anti-HRP (magenta; labels all axons) and anti-FasII (green; labels a subset of longitudinal axon pathways) (A-F) or anti-HA (G-K). Embryos carrying *elav-GAL4* alone (A) display a wild type ventral nerve cord with a ladder-like axon scaffold, two commissures per segment, and three distinct FasII-positive longitudinal pathways on either side of the midline. Misexpression of Robo1 with *elav-GAL4* (B) inhibits midline crossing, and commissures are thin or absent (arrowhead with asterisk). In embryos misexpressing mRobo1 (C) or mRobo2 (D) with *elav-GAL4*, many segments have reduced or absent commissures (arrowhead with asterisk), and in some segments FasII-positive axons ectopically cross the midline (arrow with asterisk). FasII-positive longitudinal pathways also appear disorganized in (B-D). Embryos misexpressing mRobo3.1 (E) or mRobo3.2 (F) appear wild type. (G-K) HA-tagged Robo1 and mRobos are expressed on longitudinal axons (arrows), and mRobo proteins are also detectable on midline crossing axons when present (arrowheads in H-K). (L-V) Ventral nerve cords from embryos carrying *eg-GAL4, UAS-TauMycGFP* (*TMG*), and the indicated HA-tagged *UAS-Robo* transgenes, stained with anti-HRP (magenta) and anti-GFP (green) (L-Q), or anti-HA and anti-GFP (R-V). *eg-GAL4* labels the EG and EW neurons, whose axons cross the midline in the anterior and posterior commissures, respectively. In wild type embryos (L), the EW axons cross the midline in every segment (arrowhead). Misexpression of Robo1 with *eg-GAL4* (M,R) prevents the EW axons from crossing the midline in 92.5% of segments (arrowhead with asterisk). Misexpression of mRobo1 (N,S) or mRobo2 (O,T) also prevents EW axons from crossing the midline (arrowheads with asterisks), but with lower frequency than Robo1 (82.5% for mRobo1 and 84.2% for mRobo2). EW axons cross the midline normally in *eg-GAL4/UAS-mRobo3*.*1* (P,U) and *eg-GAL4/UAS-mRobo3*.*2* (Q,V) embryos (arrowheads). Anti-HA staining in (R-V) shows that all transgenes are expressed on EW axons (arrowheads); GFP staining of the same segments is shown below for comparison.

Similarly, both mRobo1 and mRobo2 were able to strongly inhibit midline crossing of EW axons when misexpressed with *eg-GAL4*, though neither was as potent as *Drosophila* Robo1 in this assay (with 82.5% and 84.2% of segments exhibiting EW non-crossing vs 92.5%, respectively **(Figure 1L-O)**. Neither mRobo3.1 nor mRobo3.2 could repel EW axons from the midline, even though both proteins were expressed at similar levels to *Drosophila* Robo1, and clearly detectable on the axons of both EG and EW neurons, including midline crossing segments **(Figure 1P,Q)**. We conclude that mRobo1 and mRobo2 can both activate midline repulsive signaling in *Drosophila* embryonic neurons, while mRobo3.1 and mRobo3.2 appear unable to influence axon guidance of *Drosophila* neurons, whether broadly expressed in all neurons or in a more restricted subset of commissural neurons.

### Pan-neural expression of mouse Robo1 or Robo2 can partially restore midline repulsion in *Drosophila robo1* mutants

To test whether the mouse Robos could signal midline repulsion in *Drosophila* embryonic neurons in the absence of *robo1*, we performed a rescue assay by combining the *UAS-Robo1* and *UAS-mRobo* transgenes with *elav-GAL4* in *robo1* null mutant embryos. To assay midline repulsion in *robo1* mutant and rescue backgrounds, we again used anti-HRP and anti-FasII antibodies **(Figure 2)**. In wild type embryos, FasII-positive axons do not cross the midline **(Figure 2A)**, while in *robo1* mutants they cross the midline in 100% of segments **(Figure 2B)**. The high level of ectopic midline crossing in *robo1* mutants also alters the overall morphology of the axon scaffold, with the regular ladder-like arrangement of well-separated longitudinal and commissural axons partially collapsing toward the midline to produce the characteristic “roundabout” phenotype that gives *robo1* its name (Seeger et al. 1993; Kidd et al. 1998). This ectopic midline crossing can be rescued by restoring *Drosophila* Robo1 expression at high levels with *elav-GAL4* **(Figure 2C)**; expressing Robo1 at such high levels, even in embryos lacking endogenous *robo1*, also inhibits normal midline crossing and commissure formation, resulting in commissureless or nearly commissureless embryos. We found that in *robo1* mutants carrying *elav-GAL4* and *UAS-mRobo1* or *UAS-mRobo2* transgenes, some ventral nerve cord segments resembled *robo1* mutants while others exhibited ectopic midline repulsion, producing a commissureless phenotype similar to *robo1; elav-GAL4/UAS-Robo1* embryos **(Figure 2D,E)**, indicating a partial rescue of *robo1-*dependent midline repulsion. Similar to our misexpression experiments in a wild type background described above, we observed no rescue of midline repulsion or additional gain of function phenotypes in *robo1; elav-GAL4/UAS-mRobo3*.*1* or *robo1; elav-GAL4/UAS-mRobo3*.*2* embryos **(Figure 2F,G)**, and embryos of these genotypes were indistinguishable from *robo1* mutants alone.

**Figure 2.**
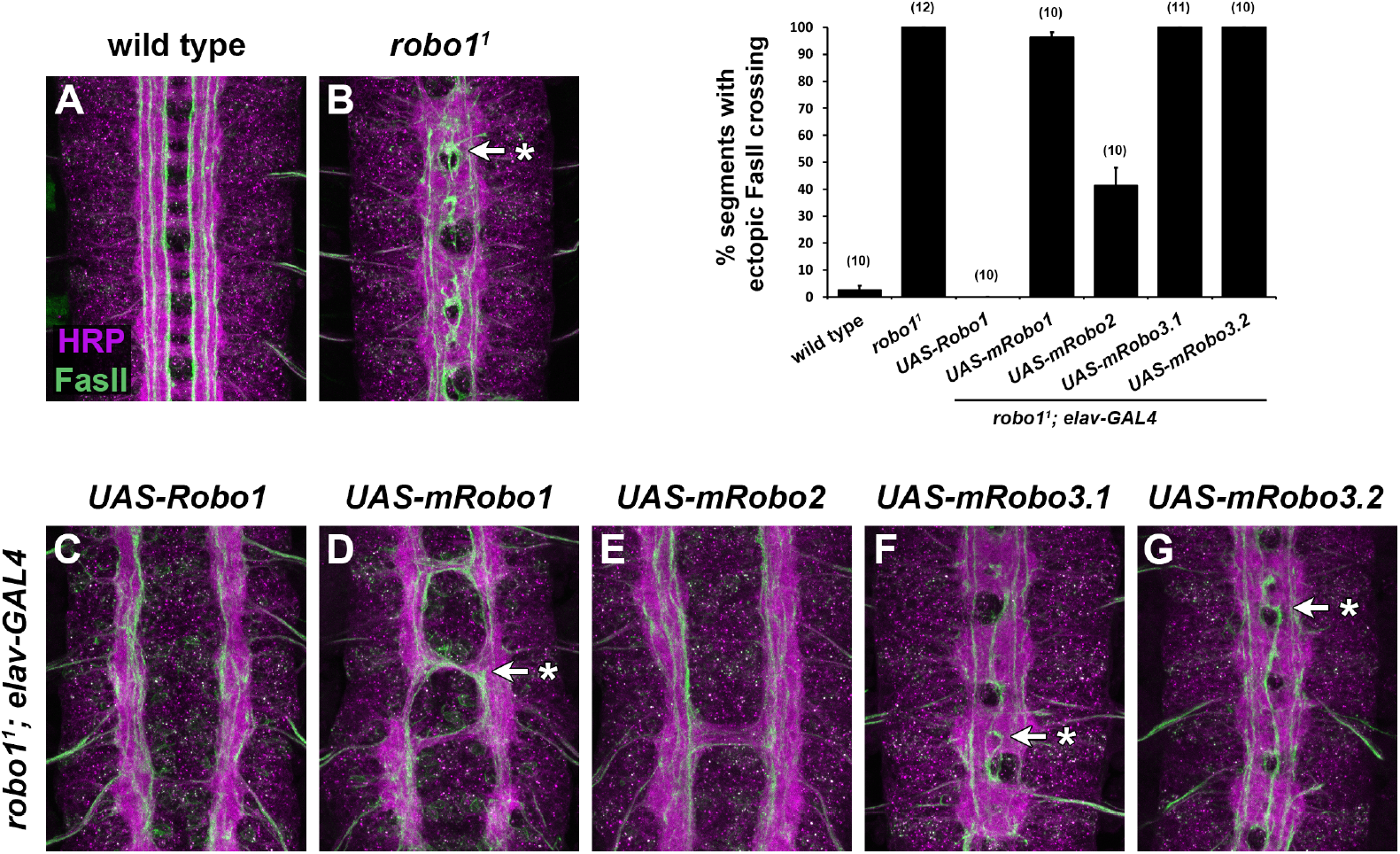
Pan-neural expression of mRobo1 or mRobo2 can partially restore midline repulsion in *Drosophila robo1* mutants. (A-G) Ventral nerve cords from *Drosophila* embryos stained with anti-HRP (magenta) and anti-FasII (green). (A) Wild type embryo; FasII-positive axon pathways do not cross the midline. (B) *robo1* mutant embryo; FasII pathways cross the midline in every segment (arrow with asterisk). (C-G) *robo1* mutant embryos carrying *elav-GAL4* and the indicated *UAS-Robo* transgenes. Pan-neural expression of *Drosophila* Robo1 (C) fully rescues the *robo1* loss of function phenotype: FasII-positive axons no longer cross the midline. Expression of mRobo1 (D) or mRobo2 (E) partially restores midline repulsion, with mRobo2 more effective than mRobo1, though FasII axons are detectable crossing the midline in both genotypes (arrows with asterisks). Expression of mRobo3.1 (F) or mRobo3.2 (G) has no effect on the *robo1* mutant phenotype. Bar graph shows quantification of ectopic midline crossing in the genotypes shown in A-G. Error bars represent s.e.m. Number of embryos scored for each genotype is indicated in parentheses.

### Expression of mouse Robos in *Drosophila* embryonic neurons via a *robo1* rescue transgene

The *GAL4/UAS-*based rescue experiments described above demonstrate that mouse Robo1 and Robo2 can partially restore midline repulsion in *robo1* mutants when expressed at high levels in all embryonic neurons. To compare the midline repulsive activity of the mouse Robos with *Drosophila* Robo1 in the context of Robo1’s endogenous expression pattern and levels, we next used a *robo1* rescue transgene to express each of the mouse Robos in the embryonic ventral nerve cord under the control of the promoter and regulatory sequences from the *Drosophila robo1* locus (Brown et al. 2015). We have previously used this transgenic approach to compare the activities of *Drosophila* Robo1 and *C*.*elegans* SAX-3 in fly embryonic neurons (Daiber et al. 2021), and to perform structure-function analyses of Robo1 ectodomain elements (Brown et al. 2015; Reichert et al. 2016; Brown et al. 2018; Brown and Evans 2020). Each of these constructs includes the endogenous signal peptide from Robo1 and a 4xHA epitope tag inserted directly upstream of the Ig1 domain, and were inserted at the same genomic location (28E7) to ensure equivalent expression levels between transgenes **(Figure 3A)**.

**Figure 3.**
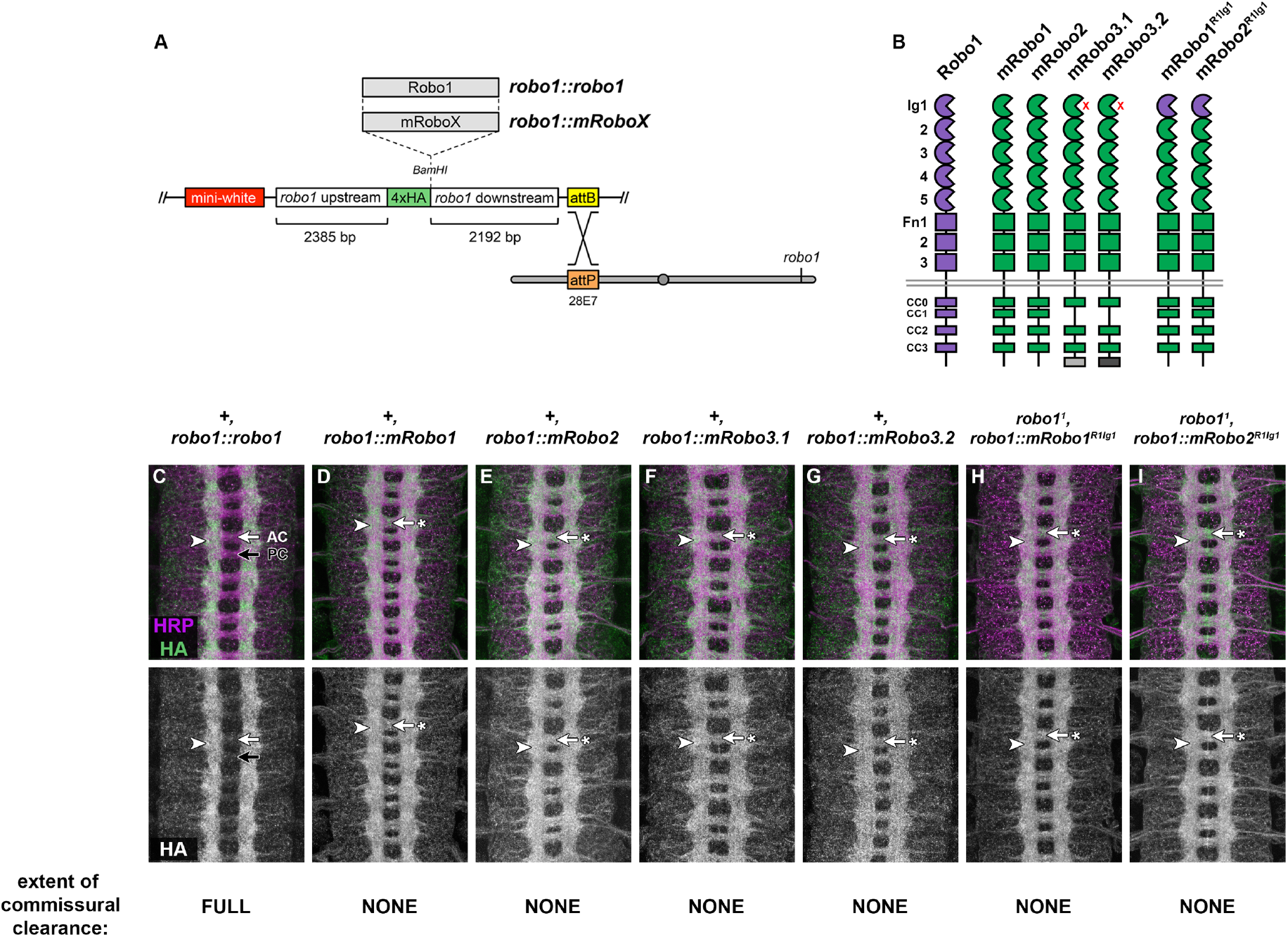
Expression and localization of mRobo and chimeric Robo1/mRobo receptors via a *robo1* genomic rescue construct. (A) Schematic of the *robo1* rescue construct (Brown et al. 2015). HA-tagged receptors are expressed under the control of regulatory regions from the *robo1* gene. All transgenes are inserted into the same genomic landing site at cytological position 28E7. (B) Schematics of *Drosophila* Robo1, mouse Robos, and Ig1 chimeric receptors. The four conserved cytoplasmic (CC) motifs present in Robo1 are conserved in mRobo1 and mRobo2; mRobo3 retains CC0, CC2, and CC3. Mouse Robo3 isoforms (mRobo3.1 and mRobo3.2) differ in an alternatively spliced exon at their C-terminus (light and dark gray boxes). Red X’s indicate lack of Slit binding of mRobo3 isoforms; all other proteins shown bind to Slit via Ig1. (C-I) Ventral nerve cords from *Drosophila* embryos stained with anti-HRP (magenta) and anti-HA (green) to reveal expression of the HA-tagged transgenic Robo proteins. HA-tagged full-length Robo1 expressed from the *robo1* rescue transgene (C) is localized to longitudinal axon pathways (arrowhead) and excluded from commissural segments in both the anterior commissure (AC, white arrow) and posterior commissure (PC, black arrow). Each of the full-length mRobo proteins (D-G) or Robo1/mRobo Ig1 chimeras (H,I) expressed from equivalent transgenes are properly localized to axons (arrowheads), but not excluded from commissures (arrows with asterisks).

When expressed from the *robo1* rescue transgene in an otherwise wild type background, HA-tagged Robo1 protein reproduces the endogenous Robo1 expression pattern, with broad expression in embryonic ventral nerve cord neurons, localization to longitudinal axons, and exclusion from commissural axon segments **(Figure 3C)** (Kidd et al. 1998; Brown et al. 2015). We found that mRobo1, mRobo2, mRobo3.1, and mRobo3.2 proteins expressed from equivalent transgenes were also properly translated, expressed at similar levels to Robo1, and localized to axons in the ventral nerve cord. However, unlike Robo1, the mouse Robo proteins were not excluded from commissures and instead were present at similar levels on longitudinal and commissural axons **(Figure 3D-G)**. We did not observe any gain of function or dominant negative-like effects in embryos carrying two copies of any of the mouse Robo transgenes in addition to two wild type copies of the endogenous *robo1* gene (e.g. *+, robo1::mRobo1*, **Figure 3D**).

### Mouse Robo receptors cannot substitute for *robo1* to signal midline repulsion at its endogenous expression levels

To determine whether mouse Robo proteins could substitute for *Drosophila* Robo1 to properly regulate midline repulsion of axons when expressed at Robo1’s endogenous expression level, we introduced our *robo1::mRobo* rescue transgenes into a *robo1* null mutant background and quantified midline repulsion using anti-FasII, which labels a subset of longitudinal axon pathways in the *Drosophila* embryonic CNS. In wild type or heterozygous *robo1/+* stage 16-17 *Drosophila* embryos, FasII-positive axons do not cross the midline **(Figure 4A)**, while they ectopically cross the midline in 100% of segments in *robo1* mutants due to an absence of *robo1-*dependent midline repulsion **(Figure 4B)**. This defect can be completely rescued by transgenic expression of *Drosophila* Robo1 using the *robo1::robo1* transgene described above **(Figure 4C)**.

**Figure 4.**
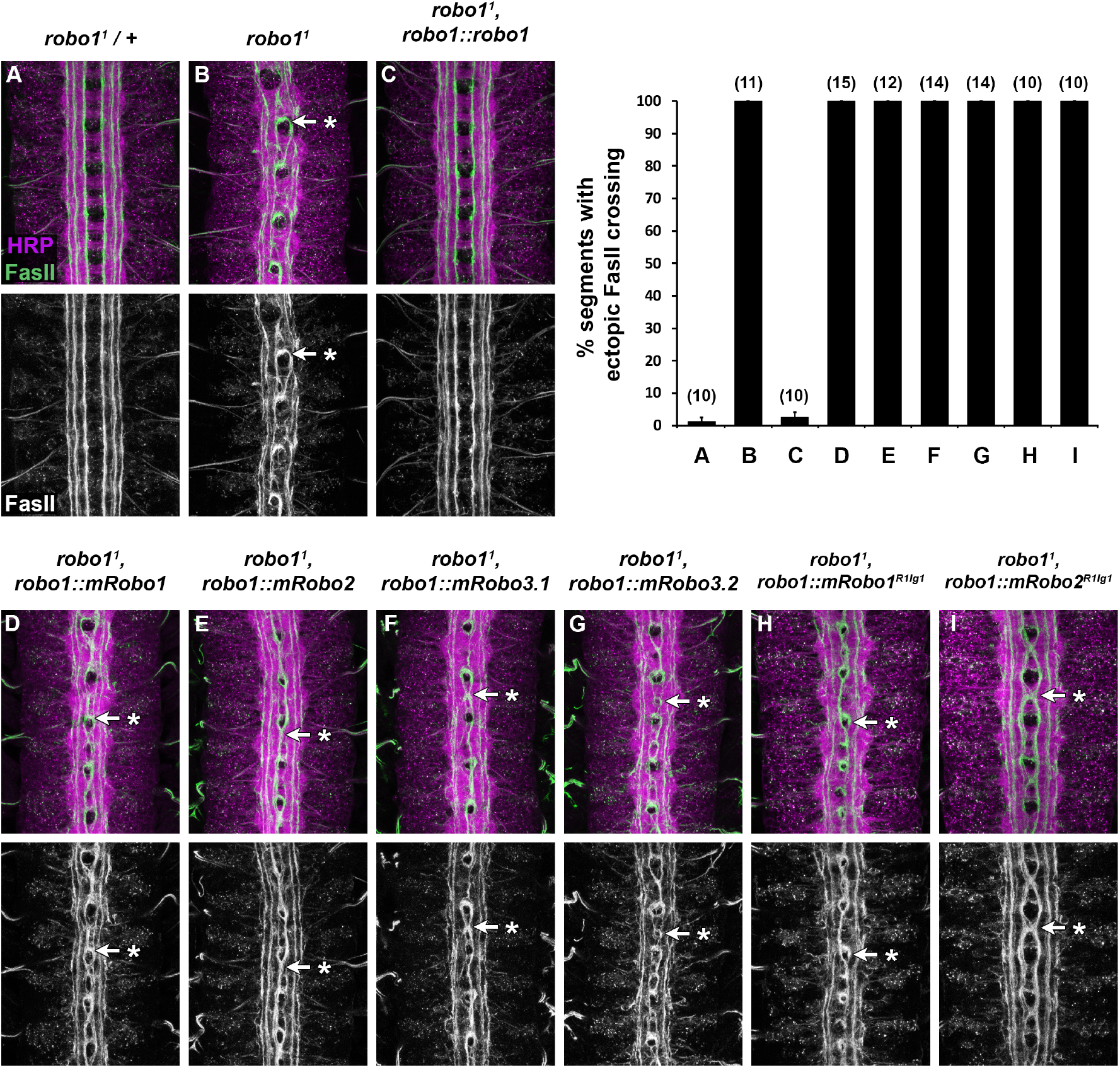
Mouse Robo receptors cannot substitute for *Drosophila* Robo1 to regulate midline crossing. (A,B) Wild type and *robo1* mutant embryonic nerve cords as in Figure 2A,B. FasII positive axons cross the midline ectopically in 100% of segments in *robo1* mutants. Restoring Robo1 expression with the *robo1::robo1* rescue transgene (C) rescues midline repulsion in *robo1* mutants completely. None of the full-length mRobo rescue transgenes (D-G) or transgenes expressing the Robo1/mRobo Ig1 chimeric receptors (H,I) can rescue ectopic FasII crossing in a *robo1* mutant background, but the overall axon scaffold appears slightly less compressed than the others in *robo1*^*1*^,*robo1::mRobo2*^*R1Ig1*^ embryos (I). Bar graph shows quantification of ectopic midline crossing in the genotypes shown in A-I. Error bars represent s.e.m. Number of embryos scored for each genotype is indicated in parentheses.

We found that none of the four mouse Robos could rescue midline repulsion in *robo1* mutants when they were expressed in the endogenous pattern and expression level of *robo1* using our rescue transgenes **(Figure 4D-G)**. In each of the four rescue backgrounds, both the degree of ectopic midline crossing of FasII-positive axons (100% of abdominal segments exhibiting ectopic crossing defects) and the overall structure of the axon scaffold was indistinguishable from *robo1* null mutants.

### Ig1 chimeric receptor variants of mouse Robo1 and Robo2

We reasoned that the decreased ability of mRobo1 and mRobo2 to substitute for Robo1 to signal midline repulsion in *Drosophila* neurons might be due to differences in response to *Drosophila* Slit. It has previously been demonstrated that *Drosophila* Slit can bind to rat Robo1 and Robo2 (rRobo1 and rRobo2) expressed on the surface of cultured cells (Brose et al. 1999), and our data presented here suggest that mouse Robo1 and Robo2 (mRobo1 and mRobo2) can detect and respond to *Drosophila* Slit in vivo, but the relative affinities of mRobo1 and mRobo2 for *Drosophila* Slit have not been directly compared to that of *Drosophila* Robo1. To test whether a simple Ig1-dependent difference in Slit affinity might account for our observed differences in receptor activity, we generated two chimeric receptors in which the Slit-binding Ig1 domains of mRobo1 and mRobo2 are replaced with Ig1 from *Drosophila* Robo1 (mRobo1^dR1Ig1^ and mRobo2^dR1Ig1^). We cloned these chimeric variant coding sequences into our *robo1* rescue construct backbone and made transgenic lines, then examined their expression and protein localization in vivo and tested each for their ability to rescue midline crossing in *robo1* mutants.

We found that exchanging the Ig1 domain of mRobo1 or mRobo2 with *Drosophila* Robo1 Ig1 did not produce any detectable alteration in the expression levels, protein stability, or axonal localization of the receptors in the embryonic CNS **(Figure 3H,I)**. The mRobo1^dR1Ig1^ and mRobo2^dR1Ig1^ chimeric receptors were not cleared from commissures, indicating that the *Drosophila* Robo1 Ig1 domain was not sufficient to confer Comm sorting sensitivity to the mouse proteins. This was not surprising, as we and others have previously shown that commissural clearance and sorting by Comm depends on the Fn3 and perimembrane regions of Robo1, not Ig1 (Gilestro 2008; Brown et al. 2018; Daiber et al. 2021). We further found that neither mRobo1^dR1Ig1^ nor mRobo2^dR1Ig1^ could restore Robo1-dependent midline repulsion in *robo1* mutants **(Figure 4H,I)**, demonstrating that exchanging the Ig1 domain (and presumably Ig1-dependent affinity for *Drosophila* Slit) was not sufficient to confer Robo1-like levels of midline repulsive signaling activity to either mRobo1 or mRobo2. However, we do note that while *robo1*^*1*^,*robo1::mRobo2*^*R1Ig1*^ embryos still exhibited ectopic FasII axon crossing in 100% of segments, the overall compression of the axon scaffold appeared qualitatively slightly less severe than in *robo1*^*1*^ mutants or *robo1*^*1*^,*robo1::mRobo2* embryos **(Figure 4I)**, suggesting a slight increase in midline repulsive signaling in mRobo2^dR1Ig1^ compared to full-length mRobo2.

## Discussion

In this paper, we have compared midline repulsive signaling by Robo family receptors in flies and mice using transgenic approaches to express *Drosophila* and mouse Robo proteins in neurons in the developing *Drosophila* embryonic CNS. We find that Robo1 and Robo2 from mouse can activate midline repulsion of axons when expressed in fly neurons, while the two reported isoforms of Robo3/Rig-1 (mRobo3.1 and mRobo3.2) have no detectable effect on midline crossing or other aspects of axon guidance when equivalently expressed. We show that mRobo1 and mRobo2 can partially restore midline repulsion in *robo1* mutant embryos when expressed at high levels via the GAL4/UAS system, but cannot rescue midline repulsion as effectively as *Drosophila* Robo1 when expressed at levels similar to endogenous Robo1. Exchanging the Slit-binding Ig1 domain of mRobo1 or mRobo2 with the equivalent domain from fly Robo1 did not confer Robo1-like levels of midline repulsive activity, suggesting that the difference in rescue ability between fly and mouse Robos cannot be explained by a simple difference in Ig1-dependent affinity for *Drosophila* Slit.

### Mouse Robos can signal midline repulsion in fly neurons

Using the GAL4/UAS system in *Drosophila* to express transgenic mouse Robo1 (Robo1) or mouse Robo2 (mRobo2) at high levels either broadly in all neurons (with *elav-GAL4*) or in a restricted subset of commissural neurons (with *eg-GAL4*), we found that mRobo1 and mRobo2 can strongly activate midline repulsive signaling and prevent midline crossing of axons in the fly embryonic CNS, displaying gain of function phenotypes that were nearly as strong as that of *Drosophila* Robo1.

In contrast, neither reported isoform of mouse Robo3/Rig-1 (mRobo3.1 or mRobo3.2) (Chen et al. 2008) had any detectable effect on midline crossing when expressed under similar conditions. Importantly, mammalian Robo3 homologs (including mRobo3) are not Slit receptors and instead respond to the novel ligand NELL2 (Zelina et al. 2014; Jaworski et al. 2015), which does not appear to have a *Drosophila* ortholog. This functional divergence is a likely explanation for the absence of any gain of function phenotypes when mRobo3.1 or mRobo3.2 are expressed in *Drosophila* neurons.

All three mouse Robo receptors share a similar ectodomain structure with *Drosophila* Robo1 (5 Ig + 3 Fn), while the cytoplasmic domain sequences are more divergent. The four conserved cytoplasmic (CC) motifs present in *Drosophila* Robo1 are also found in mouse Robo1 and Robo2 (indeed, CC1-CC3 were originally defined based on their conservation in *Drosophila* Robo1 and human Robo1). Our observation that mRobo1 and mRobo2 can apparently respond to fly Slit is consistent with previous studies showing that *Drosophila* Slit can bind to mammalian Robo receptors in cultured cells (Brose et al. 1999), and suggests that both mouse receptors are capable of interacting with and activating downstream components of the Slit-Robo signaling pathway. However, it is important to note that we cannot formally rule out the possibility that the midline repulsion we observe under conditions of mRobo misexpression could instead be caused by heteromultimer formation with *Drosophila* Robo1 (or, in our rescue experiments in a *robo1* mutant background, with *Drosophila* Robo2 or Robo3) rather than direct binding to *Drosophila* Slit or direct interaction with downstream cytoplasmic effectors.

### Rescue of *robo1-*dependent midline repulsion by mouse Robos

In addition to the gain of function experiments carried out in a wild type background (i.e. in embryos where endogenous Robo1 expression was present), we found that mRobo1 and mRobo2 could each restore some degree of midline repulsion in *Drosophila* embryos lacking endogenous *robo1*, although neither receptor was as effective at rescue as *Drosophila* Robo1 itself. When expressed at levels equivalent to endogenous *robo1* using a transgenic rescue construct, neither mRobo1 nor mRobo2 exhibited any detectable rescue of the ectopic midline crossing caused by loss of *robo1*. However, both mRobo1 and mRobo2 could partially rescue the *robo1* mutant phenotype when expressed at high levels in embryonic neurons using *elav-GAL4*. Notably, the extent of rescue we observed with mRobo2 under these conditions was greater than that conferred by *Drosophila* Robo2 or Robo3 under equivalent conditions (Evans and Bashaw 2012), supporting the idea that midline repulsive signaling by *Drosophila* Robo1 is more similar to mRobo1 and mRobo2 than its *Drosophila* paralogs.

We have previously reported a similar series of experiments examining the midline repulsive activity of the single *C. elegans* Robo family receptor (SAX-3) in *Drosophila* neurons (Daiber et al. 2021). We note some interesting contrasts between the results of those experiments and the present study. For example, we observed no rescue in *robo1* mutants expressing SAX-3 via *GAL4/UAS*, but did see partial rescue at endogenous *robo1* expression levels using our *robo1* rescue transgene, while with mRobo1/mRobo2 we saw partial rescue with *GAL4/UAS* but no rescue with the *robo1* rescue transgene. In addition, we observed severe disorganization of the axon scaffold when expressing SAX-3 via *elav-GAL4* in a *robo1* mutant background, but saw no such effects with mRobo1 or mRobo2. Finally, expression of mRobo1 or mRobo2 via our rescue transgene in a wild type background (i.e. with two functional copies of *robo1*) did not cause any detectable dominant negative-like effects, but expression of SAX-3 in the same context did induce a mild dominant negative effect (Daiber et al. 2021).

Justice and colleagues have reported that pan-neural expression of human Robo1 (hRobo1) in the *Drosophila* embryonic nerve cord via *scabrous-GAL4* causes a subtle increase in midline crossing, presumably via a dominant negative-like effect on *Drosophila* Robo1 (Justice et al. 2017). This contrasts with our observations, where we saw no apparent dominant-negative effect of expressing mRobo1 or mRobo2 under any conditions. However, these experiments may not be directly comparable, as each involves expression of Robo homologs from different vertebrate species (human vs mouse) using different GAL4 lines (*sca-GAL4* vs *elav-GAL4)* and responder lines with different numbers of UAS repeats (*5xUAS-hRobo1* vs *10xUAS-mRobo1* and *10xUAS-mRobo2*) likely producing differences in the timing, pattern, and/or levels of expression in the different experiments.

### Differences in Slit affinity vs differences in receptor activation/signaling mechanism(s)

Slit binding by Robo receptors in mammals and flies appears to depend primarily on the N-terminal Ig1 domain. We considered the possibility that species-specific differences in Slit affinity (i.e. higher affinity of *Drosophila* Robo1 for *Drosophila* Slit compared to mRobo1’s or mRobo2’s affinity for *Drosophila* Slit) might account for the difference in effectiveness of midline repulsive signaling between fly Robo1 and the two mRobos. We reasoned that replacing Ig1 of mRobo1 or mRobo2 with the Ig1 domain of *Drosophila* Robo1 might increase their affinity for *Drosophila* Slit and thus confer Robo1-equivalent midline repulsive activity. However, neither mRobo1^R1Ig1^ nor mRobo2^R1Ig1^ could rescue midline repulsion as effectively as *Drosophila* Robo1, suggesting that a simple Ig1-dependent Slit affinity difference cannot account for the observed differences in midline repulsive activity.

An alternative possibility is that perhaps ectodomain-dependent differences in signaling mechanism (conformational change, etc) could account for this difference. We note that previous in vitro structural studies of human Robo1 and Robo2 suggest functional roles of structural domains other than Ig1 in these receptors, for example Ig3/D3- and Ig4/D4-dependent dimerization (Aleksandrova et al. 2017 Dec 28; Barak et al. 2019) that appear to be conserved and necessary for signaling by *C. elegans* SAX-3 (Barak et al. 2019). In contrast, we have previously shown that Ig3 and Ig4 are dispensable for in vivo midline repulsive signaling by *Drosophila* Robo1 (indeed the entire Ig2-Ig5 region can be deleted without any detectable effect on midline repulsion) (Reichert et al. 2016; Brown and Evans 2020), perhaps indicating that activation of *Drosophila* Robo1 may involve a different structural mechanism than activation of mammalian Robo1/Robo2 and *C. elegans* SAX-3. Whether or not this possible difference in receptor activation might account for the inability of mRobo1, mRobo2, and SAX-3 to fully rescue Robo1-dependent midline repulsive signaling in *Drosophila* will require further comparative studies to answer.

## Acknowledgments

We thank Mike Fleming and Wenqin Luo for providing mouse embryonic cDNA. Stocks obtained from the Bloomington Drosophila Stock Center [National Institutes of Health (NIH) grant P40 OD-018537] were used in this study. Monoclonal antibodies were obtained from the Developmental Studies Hybridoma Bank, created by the Eunice Kennedy Shriver National Institute of Child Health and Human Development of the NIH and maintained at The Department of Biology, University of Iowa, Iowa City, IA 52242. This work was supported by NIH grant R15 NS-098406 (T.A.E.).

## References

Aleksandrova N, Gutsche I, Kandiah E, Avilov SV, Petoukhov MV, Seiradake E, McCarthy AA. 2017 Dec 28. Robo1 Forms a Compact Dimer-of-Dimers Assembly. Structure. doi:10.1016/j.str.2017.12.003. http://linkinghub.elsevier.com/retrieve/pii/S0969212617304021.

Barak R, Yom-Tov G, Guez-Haddad J, Gasri-Plotnitsky L, Maimon R, Cohen-Berkman M, McCarthy AA, Perlson E, Henis-Korenblit S, Isupov MN, et al. 2019. Structural Principles in Robo Activation and Auto-inhibition. Cell. 177(2):272-285.e16. doi:10.1016/j.cell.2019.02.004.

Bashaw GJ, Kidd T, Murray D, Pawson T, Goodman CS. 2000. Repulsive axon guidance: Abelson and Enabled play opposing roles downstream of the roundabout receptor. Cell. 101(7):703–715.

Brose K, Bland KS, Wang KH, Arnott D, Henzel W, Goodman CS, Tessier-Lavigne M, Kidd T. 1999. Slit proteins bind Robo receptors and have an evolutionarily conserved role in repulsive axon guidance. Cell. 96(6):795–806.

Brown HE, Evans TA. 2020. Minimal structural elements required for midline repulsive signaling and regulation of Drosophila Robo1. PLoS ONE. 15(10):e0241150. doi:10.1371/journal.pone.0241150.

Brown HE, Reichert MC, Evans TA. 2015. Slit Binding via the Ig1 Domain Is Essential for Midline Repulsion by Drosophila Robo1 but Dispensable for Receptor Expression, Localization, and Regulation in Vivo. G3: Genes|Genomes|Genetics. 5(11):2429–2439. doi:10.1534/g3.115.022327.

Brown HE, Reichert MC, Evans TA. 2018. In Vivo Functional Analysis of Drosophila Robo1 Fibronectin Type-III Repeats. G3: Genes|Genomes|Genetics. 8(2):621–630. doi:10.1534/g3.117.300418.

Carranza A, Howard LJ, Brown HE, Ametepe AS, Evans TA. 2023. Slit-independent guidance of longitudinal axons by Drosophila Robo3. bioRxiv. doi:10.1101/2023.05.08.539901. [accessed 2023 Jun 8]. http://biorxiv.org/lookup/doi/10.1101/2023.05.08.539901.

Chen Z, Gore BB, Long H, Ma L, Tessier-Lavigne M. 2008. Alternative splicing of the Robo3 axon guidance receptor governs the midline switch from attraction to repulsion. Neuron. 58(3):325–332. doi:10.1016/j.neuron.2008.02.016.

Daiber T, VanderZwan-Butler CJ, Bashaw GJ, Evans TA. 2021. Conserved and divergent aspects of Robo receptor signaling and regulation between Drosophila Robo1 and C. elegans SAX-3. Genetics. 217(3). doi:10.1093/genetics/iyab018. http://eutils.ncbi.nlm.nih.gov/entrez/eutils/elink.fcgi?dbfrom=pubmed&id=33789352&retmode=ref&cmd=prlinks.

Dittrich R, Bossing T, Gould AP, Technau GM, Urban J. 1997. The differentiation of the serotonergic neurons in the Drosophila ventral nerve cord depends on the combined function of the zinc finger proteins Eagle and Huckebein. Development. 124(13):2515– 2525.

Evans TA, Bashaw GJ. 2010. Functional diversity of Robo receptor immunoglobulin domains promotes distinct axon guidance decisions. Curr Biol. 20(6):567–572. doi:10.1016/j.cub.2010.02.021.

Evans TA, Bashaw GJ. 2012. Slit/Robo-mediated axon guidance in Tribolium and Drosophila: Divergent genetic programs build insect nervous systems. Dev Biol. 363(1):266–278. doi:10.1016/j.ydbio.2011.12.046.

Evans TA, Santiago C, Arbeille E, Bashaw GJ. 2015. Robo2 acts in trans to inhibit Slit-Robo1 repulsion in pre-crossing commissural axons. Elife. 4:e08407. doi:10.7554/eLife.08407.

Gilestro GF. 2008. Redundant mechanisms for regulation of midline crossing in Drosophila. PLoS ONE. 3(11):e3798. doi:10.1371/journal.pone.0003798.

Hauptman G, Reichert MC, Abdal Rhida MA, Evans TA. 2022. Characterization of enhancer fragments in Drosophila robo2. Fly. 16(1):312–346. doi:10.1080/19336934.2022.2126259.

Jaworski A, Long H, Tessier-Lavigne M. 2010. Collaborative and specialized functions of robo1 and robo2 in spinal commissural axon guidance. J Neurosci. 30(28):9445–9453. doi:10.1523/JNEUROSCI.6290-09.2010.

Jaworski A, Tom I, Tong RK, Gildea HK, Koch AW, Gonzalez LC, Tessier-Lavigne M. 2015. Operational redundancy in axon guidance through the multifunctional receptor Robo3 and its ligand NELL2. Science. 350(6263):961–965. doi:10.1126/science.aad2615.

Justice ED, Barnum SJ, Kidd T. 2017. The WAGR syndrome gene PRRG4 is a functional homologue of the commissureless axon guidance gene. PLoS Genet. 13(8):e1006865. doi:10.1371/journal.pgen.1006865.

Kidd T, Brose K, Mitchell KJ, Fetter RD, Tessier-Lavigne M, Goodman CS, Tear G. 1998. Roundabout controls axon crossing of the CNS midline and defines a novel subfamily of evolutionarily conserved guidance receptors. Cell. 92(2):205–215.

Kidd T, Evans T. 2023a. Collection, Fixation, and Antibody Staining of Drosophila Embryos. Cold Spring Harbor Protocols.

Kidd T, Evans T. 2023b. Ventral Nerve Cord Dissection and Microscopy of Drosophila Embryos. Cold Spring Harbor Protocols.

Long H, Sabatier C, Ma L, Plump AS, Yuan W, Ornitz DM, Tamada A, Murakami F, Goodman CS, Tessier-Lavigne M. 2004. Conserved roles for Slit and Robo proteins in midline commissural axon guidance. Neuron. 42(2):213–223.

O’Donnell MP, Bashaw GJ. 2013. Src Inhibits Midline Axon Crossing Independent of Frazzled/Deleted in Colorectal Carcinoma (DCC) Receptor Tyrosine Phosphorylation. J Neurosci. 33(1):305–314. doi:10.1523/JNEUROSCI.2756-12.2013.

Ogienko AA, Andreyeva EN, Omelina ES, Oshchepkova AL, Pindyurin AV. 2020. Molecular and cytological analysis of widely-used Gal4 driver lines for Drosophila neurobiology. BMC Genet. 21(S1):96. doi:10.1186/s12863-020-00895-7.

Pak JS, DeLoughery ZJ, Wang J, Acharya N, Park Y, Jaworski A, Özkan E. 2020. NELL2-Robo3 complex structure reveals mechanisms of receptor activation for axon guidance. Nat Commun. 11(1):1489–14. doi:10.1038/s41467-020-15211-1.

Pappu KS, Morey M, Nern A, Spitzweck B, Dickson BJ, Zipursky SL. 2011. Robo-3–mediated repulsive interactions guide R8 axons during Drosophila visual system development. Proceedings of the National Academy of Sciences. 108(18):7571.

Patel NH. 1994. Imaging neuronal subsets and other cell types in whole-mount Drosophila embryos and larvae using antibody probes. Methods in cell biology. 44:445–487.

Rajagopalan S, Nicolas E, Vivancos V, Berger J, Dickson BJ. 2000. Crossing the midline: roles and regulation of Robo receptors. Neuron. 28(3):767–777.

Rajagopalan S, Vivancos V, Nicolas E, Dickson BJ. 2000. Selecting a longitudinal pathway: Robo receptors specify the lateral position of axons in the Drosophila CNS. Cell. 103(7):1033–1045.

Reichert MC, Brown HE, Evans TA. 2016. In vivo functional analysis of Drosophila Robo1 immunoglobulin-like domains. Neural development. 11(1):15. doi:10.1186/s13064-016-0071-0.

Sabatier C, Plump AS, Ma L, Brose K, Tamada A, Murakami F, Lee EY-HP, Tessier-Lavigne M. The divergent Robo family protein Rig-1/Robo3 is a negative regulator of slit responsiveness required for midline crossing by commissural axons. Cell. 117(2):157– 169.

Santiago C, Labrador J-P, Bashaw GJ. 2014. The homeodomain transcription factor Hb9 controls axon guidance in Drosophila through the regulation of Robo receptors. Cell Rep. 7(1):153–165. doi:10.1016/j.celrep.2014.02.037.

Schindelin J, Arganda-Carreras I, Frise E, Kaynig V, Longair M, Pietzsch T, Preibisch S, Rueden C, Saalfeld S, Schmid B, et al. 2012. Fiji: an open-source platform for biological-image analysis. Nat Methods. 9(7):676–682. doi:10.1038/nmeth.2019.

Seeger M, Tear G, Ferres-Marco D, Goodman CS. 1993. Mutations affecting growth cone guidance in Drosophila: genes necessary for guidance toward or away from the midline. Neuron. 10(3):409–426.

Simpson JH, Bland KS, Fetter RD, Goodman CS. 2000. Short-range and long-range guidance by Slit and its Robo receptors: a combinatorial code of Robo receptors controls lateral position. Cell. 103(7):1019–1032.

Simpson JH, Kidd T, Bland KS, Goodman CS. 2000. Short-range and long-range guidance by slit and its Robo receptors. Robo and Robo2 play distinct roles in midline guidance. Neuron. 28(3):753–766.

Spitzweck B, Brankatschk M, Dickson BJ. 2010. Distinct Protein Domains and Expression Patterns Confer Divergent Axon Guidance Functions for Drosophila Robo Receptors. Cell. 140(3):409–420. doi:10.1016/j.cell.2010.01.002.

Zallen JA, Yi BA, Bargmann CI. 1998. The conserved immunoglobulin superfamily member SAX-3/Robo directs multiple aspects of axon guidance in C. elegans. Cell. 92(2):217– 227.

Zelina P, Blockus H, Zagar Y, Péres A, Friocourt F, Wu Z, Rama N, Fouquet C, Hohenester E, Tessier-Lavigne M, et al. 2014. Signaling switch of the axon guidance receptor Robo3 during vertebrate evolution. Neuron. 84(6):1258–1272. doi:10.1016/j.neuron.2014.11.004.

Zlatic M, Landgraf M, Bate M. 2003. Genetic specification of axonal arbors: atonal regulates robo3 to position terminal branches in the Drosophila nervous system. Neuron. 37(1):41–51.

